# Identifying the factors governing internal state switches during nonstationary sensory decision-making

**DOI:** 10.1101/2024.02.02.578482

**Authors:** Zeinab Mohammadi, Zoe C. Ashwood, The International Brain Laboratory, Jonathan W. Pillow

## Abstract

Recent work has revealed that mice do not rely on a stable strategy during perceptual decision-making, but switch between multiple strategies within a single session [1, 2]. However, this switching behavior has not yet been characterized in non-stationary environments, and the factors that govern switching remain unknown. Here we address these questions using an internal state model with input-driven transitions. Our approach relies on a hidden Markov model (HMM) with two sets of per-state generalized linear models (GLMs): a set of Bernoulli GLMs for modeling the animal’s state- and stimulus-dependent choice on each trial, and a multinomial GLM for modeling input-dependent transitions between states.

We used this model to analyze a dataset from the International Brain Laboratory (IBL), in which mice performed a binary decision-making task with non-stationary stimulus statistics. We found that mouse behavior in this task was accurately described by a four-state model. This model contained two “engaged” states, in which performance was good despite slight left and right biases, and two “disengaged” states, where performance was low and exhibited with larger left and right biases, respectively. Our analyses revealed that mice preferentially used left-bias strategies during left-bias stimulus blocks, and right-bias strategies during right-bias stimulus blocks, meaning that they could achieve reasonably high performance even in disengaged states simply by biasing choice toward the side with greater prior probability. Our model showed that past choices and past stimuli predicted transitions between left- and right-bias states, while past rewards predicted transitions between engaged and disengaged states. In particular, greater past reward predicted transition to disengaged states, suggesting that disengagement may be associated with satiety.

## 1 Introduction

A large literature has sought to characterize the computational mechanisms governing perceptual decision-making [3–8]. Recent work has revealed that these computations are generally not constant from one trial to another, even within a single session. Rather, animals frequently alternate between different strategies for mapping sensory evidence to decisions. [1, 2, 9–13]. However, the factors governing these state switches remains an important open problem.

Hidden Markov Models (HMMs) provide a powerful statistical framework for modeling state-dependent behavior [14, 15]. In particular, a Hidden Markov Model (HMM) with Generalized Linear Model observations (GLM) provides a modeling framework (GLM-HMM) that has recently been used to identify latent internal states from decision-making data. These states correspond to distinct decision-making strategies, each parameterized by a set of GLM weights that describe how the animal integrates covariates to make decisions in that particular state (see Methods).

However, previous work relied on HMMs with fixed transition probabilities between states, and thus provided no insight into the factors that govern switching between various strategies. Consequently, a gap exists in approaches that can track and model the transition patterns in internal states while substantiating the impact of latent states on animal behavior. To address this, a statistical model incorporating time-varying transition probabilities between states is imperative to capture the dynamic nature of the animal brain. The resulting model is a HMM with two sets of per-state GLMs. These GLMs are referred to as GLM-O (observation GLM) for observation and GLM-T (transition GLM) for transition. GLM-O calculates the output probability for each state, while GLM-T determines the transition probabilities to subsequent states.

We applied our model to analyze decision-making data sourced from a foraging-like task [16]. The stimulus statistics change in blocks, making it a task with a richer temporal structure than the one considered in [1]. Therefore, it would be optimal and natural to change states in response to changes in the stimulus statistics. We conducted a comparison of various GLM-HMM configurations, each featuring a different number of states. Our model selection incorporated a cross-validation technique, aimed at identifying the model that could capture the subtleties within the data most effectively. After an evaluation of log-likelihood values, the GLM-HMM with four distinct states, namely “Engaged-L”, “Engaged-R” “Biased-L” and “Biased-R” outperformed the 1-state model by a considerable margin. Notably, this four-state model also proved to be more interpretable, making it an ideal choice for this study.

Our analyses reveal that this four-state model provides a more robust explanation for the underlying data structure. In this model, while the stimulus plays a significant role, states characterized by a bias weight effectively capture the nuances within biased data blocks. This achievement owes much to the transition GLM, which adeptly captures transition patterns between states, and the observation GLM, which accurately depicts the animal’s choices.

In summary, our study underscores the effectiveness of the HMM incorporating two sets of Bernoulli GLM and multinomial GLM in representing animal decision behavior and the transition probabilities between states. This approach is particularly valuable in scenarios where the probability of stimulus appearing on different sides of the screen is variable and frequently changes during a subset of trials.

## 2 Results

### Modeling internal state transitions underlying decision-making

To model animal decision-making behavior, we used an internal state model known as the GLM-HMM [1, 2, 9]. The basic GLM-HMM consists of a Hidden Markov Model (HMM) in which each state corresponds to a decision-making strategy parameterized by a Bernoulli generalized linear model (GLM). Each GLM is effectively a logistic regression model with a set of weights describing how the animal integrates external inputs (e.g., sensory stimulus, bias, trial history) to make decisions in that particular state.

This methodology introduces an HMM parameterized by two distinct sets of GLMs, enabling the effective capture of time-variation model characteristics through non-constant between-states transition probabilities. So, HMM serves as a temporal probabilistic model that characterizes states through a discrete random variable. Within this model, diverse animal decision-making strategies are encapsulated by hidden states, each corresponding to a specific GLM regression approach. The proposed input-output GLM-HMM encompasses two categories of GLMs termed Observational GLM (GLM-O) and Transitional GLM (GLM-T). The former calculates the output probability for each state, representing the animal’s choice, while the latter generates the transition probabilities to different states. Therefore, GLM-O delineates the influence of various regressors on animal choice within each state, while GLM-T quantifies the impact of covariates on state transition probabilities. These two GLMs are endowed with distinct regression weight sets. The design of this GLM-HMM construct facilitates the calculation of transition probabilities to other states at each instance, relying on sensory input, previous reward, and other regressors to formulate the transition matrix at the current state. Consequently, based on the derived outcomes, it effectively predicts the subsequent state of the animal. Importantly, this modeling paradigm allows for incorporating an arbitrary number of states and their modulation through diverse transition and observation regressors.

While the mathematical formulation of the GLM presents the output as a linear combination of diverse covariates, here manifested as the animal’s choice, it is more precise to characterize the animal’s decision-making behavior with a time-dependent latent state model. This motivated the adoption of the GLM-HMM, where each state features a distinct set of GLM weights, thereby introducing the flexibility of temporal dependency within each animal strategy. In contrast to previous models [1, 2], where probabilistic transitions between states relied on fixed probabilities in a transition matrix, our approach incorporates time-dependent transitions (see Method section).

Within this model, an array named “initial states values” is introduced, with the constraint that the sum of its elements equals 1. The latent model comprises 2*K* independent GLMs (with *K* as the number of states) encompassing *K* Bernoulli GLMs for GLM-O and *K* multinomial GLMs for GLM-T. These GLMs are associated with two sets of parameters: observation weights and transition weights. Weight vectors specific to each state elucidate the impact of regressors/inputs on the output. The outputs for observation weights and transition weights correspond to mice choices and transition probabilities to subsequent states, respectively.

Therefore, in this context, the observation GLM can be described as:

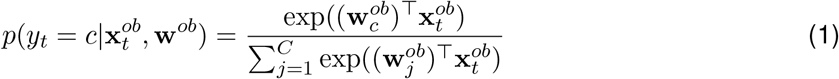

Here, 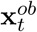 denotes the observation covariates at trial *t*. Additionally, 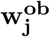 corresponds to the observation weights associated with the j-th outcome, and *C* is the number of possible outcomes. The variable *y*_*t*_ presents the animal’s choice at trial *t*. A value of 1 for *y*_*t*_ indicates the mouse turning the wheel to the right in the IBL task [16], which will be described in detail in the next section, while 0 indicates a turn to the left. Therefore, in this case, *C* equals 2 and we can write the above equation as:

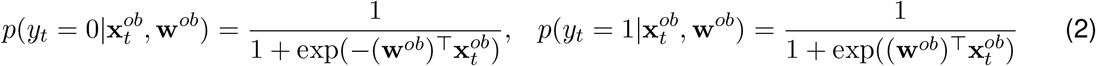

In the context of a multinomial GLM, for the transition model, the GLM output reflects different probabilities for transitions between states. In this case, 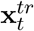 presents the transition covariates at trial *t* and 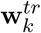 is the transition weights associated with state *k*. So for the transition probability to state *k* at trial *t*, denoted as *z*_*t*_, we can write:

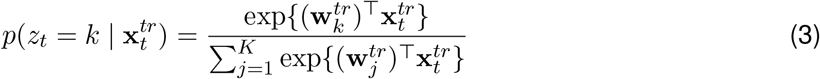

Here, *K* represents the total number of states.

### An analytical framework for modeling mice data in decision-making task

We fit the described GLM-HMM to a decision-making task data called IBL dataset [16]. This data is gathered from perceptual decision-making experiments conducted in several laboratories. In this task, mice detect the direction of a Gabor patch on the screen and subsequently turn the wheel to the right or left to indicate the stimulus direction [17]. We have used this IBL data, 37 mice, to fit the GLM-HMM framework with a different number of states. Our animal selection criteria involved including those with a minimum of 30 sessions and incorporating sessions characterized by a low number of error trials, where the animal either did not make a choice or timed out.

In fitting the model, Maximum A Posteriori (MAP) was employed. The model parameters, denoted as Θ = {**w**^*tr*^, **w**^*ob*^, ***π***}, include the transition weights, observation weights and the initial state distribution respectively for all states.

In this fitting procedure, observations and a set of latent states, denoted as **Z** = *z*_1_, …, *z*_*T*_ and **Y** = *y*_1_, …, *y*_*T*_ respectively, coexist with two defined sets of covariates: 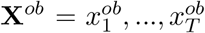 representing observation covariates and 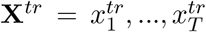 representing transition covariates. During each trial, we utilize the specified GLM-HMM parameters to compute a joint probability distribution that encompasses both the states and the animals’ decisions. Subsequently, we evaluate the model’s log-likelihood based on this calculated joint probability distribution. This relationship can be written as:

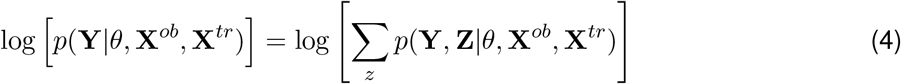

Fig. 1a shows an illustrative Diagram of the IBL sensory decision-making task. Each experimental trial encompasses the presentation of a sinusoidal grating, characterized by gradient values ranging from 0% to 100%. This grating stimulus is selectively presented on either the left or right periphery of the visual display. Subsequently, mice are mandated to discriminate the spatial location of the grating and communicate their decision via rotational manipulation of a wheel, resulting in a left or right turn, which corresponds to the perceived location of the grating stimulus. Successful execution of this task merits a water reward. For further insights into this experiment, please refer to the detailed description of the IBL task [16, 17].

**Figure 1:**
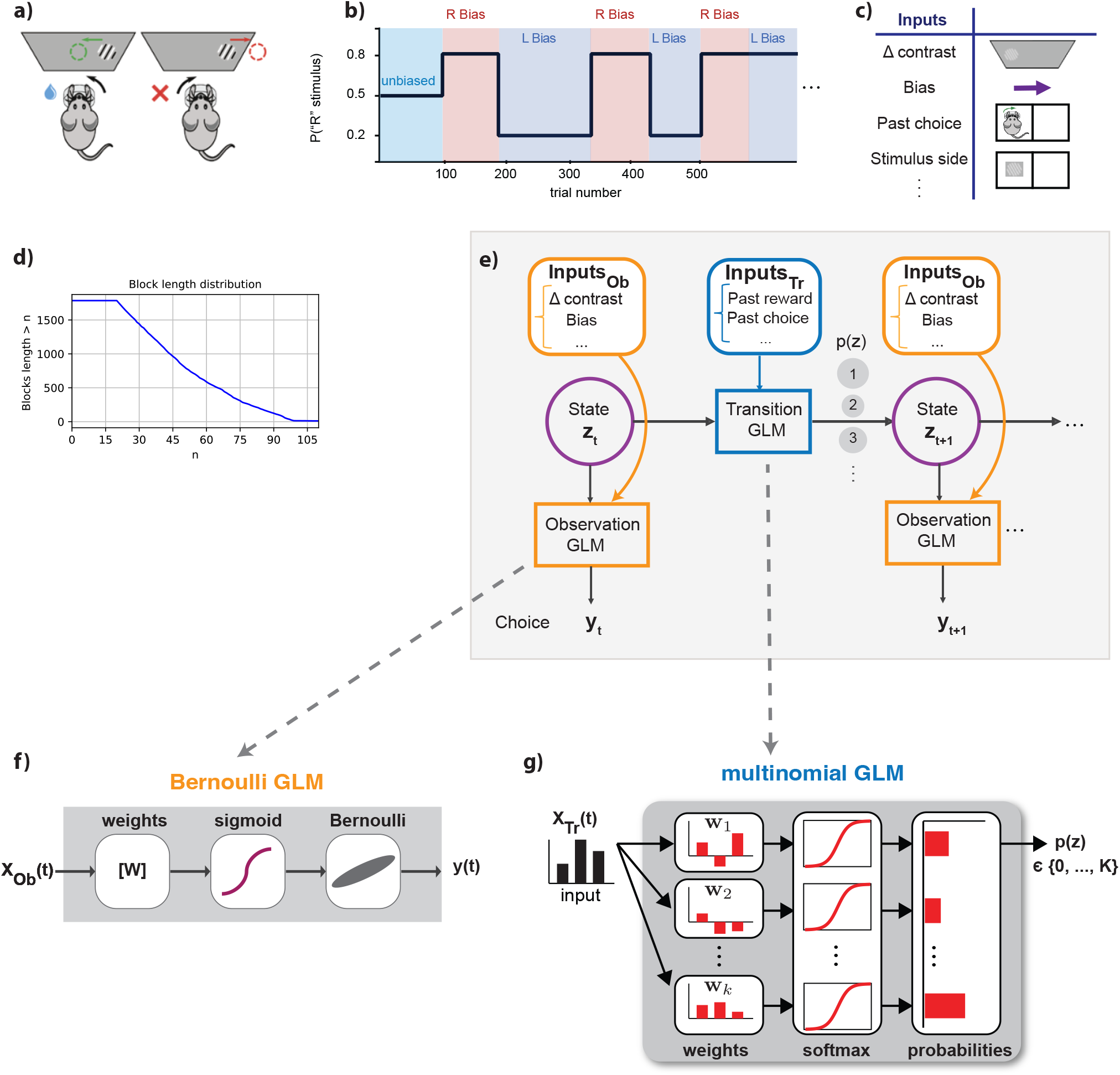
GLM-HMM with Bernoulli and multinomial GLMs for the decision-Making task. **(a)** IBL sensory decision-making paradigm; experimental trials involve showing mice a sinusoidal grating with varying contrast (0%-100%) on either the left or right side of a screen. Mice determine the grating’s location and convey their choice by turning a wheel left or right, earning a water reward for accurate responses [16]. **(b)** Stimulus structure of the IBL task; biased and unbiased blocks were present after training. This diagram shows that sessions commenced with unbiased trials with a 50:50 probability and transitioned to variable-length biased blocks with 20:80 or 80:20 ratios for the right or left sides. **(c)** Diverse inputs contribute to the composite structure of the GLM-HMM. This framework encompasses two distinct sets of inputs: transition inputs (*X*_*T r*_) and observation inputs (*X*_*Ob*_). We present a subset of these inputs herein, aimed at exemplifying the variability within these categories. **(d)** Depicting the count of biased data blocks with lengths exceeding a specified threshold value (denoted as ‘n’). **(e)** Model diagram (in a box with a light gray background color): Incorporating the GLM-HMM paradigm featuring distinct GLMs tailored for transitions and observations. Each state within this framework integrates a state-specific Bernoulli GLM (GLM-O) and a multinomial GLM (GLM-T), respectively accounting for choice probability and state transitions probability. **(f)** Bernoulli GLM to model binary observations (mouse choices) while considering relevant covariates. **(g)** Multinomial GLM mapping covariate effects to represent the transitions probabilities between different states.

In the IBL task, after training mice in the foundational purely sensory task, they were introduced to an advanced paradigm where optimal performance necessitated the fusion of sensory perception with recent experience. Specifically, block-wise biases were incorporated into the probability distribution of stimulus locations, thereby influencing the more probable correct choice. Each session commenced with an unbiased trials block, offering an equal 50:50 probability of left versus right stimulus locations. The length of the unbiased block was 90 trials for all sessions of the task (Fig. 1b). Subsequent trial blocks alternated variably and exhibited biases toward the right and left. The probability distribution skewed at a 20:80 and an 80:20 ratio for the right and left sides, respectively. The length of these biased blocks ranged from 20 to 100 trials. The transition between these biased blocks was not overtly indicated, necessitating the mice to extrapolate a prior estimation for stimulus location based on recent task statistics. This intricate task compels the mice to assimilate information across multiple trials, strategically employing their prior knowledge to inform their perceptual decisions.

It should be noted that our primary focus here is on designing the transitional and observational GLMs for a GLM-HMM to fit the experimental data appropriately, incorporating relevant weights for accurate state prediction. For each section of the data, comprising trials or sessions, the model utilizes the stimulus and other covariates to calculate the transitional probabilities of the mice internal state at each time bin (see Methods section). We employ this model to gain insights into the mice strategies within each latent state as they transition to other states.

Within this modeling framework, two distinct sets of inputs were incorporated, encompassing transition and observation inputs. In this context, the observed output corresponded to the animal’s decision, manifesting as a binary value indicative of the direction in which the wheel was turned—either right or left. A multitude of covariates were additionally integrated into the model and will be elucidated herein. Specifically, the “stimulus” parameter was defined as the contrast level of a sinusoidal grating, spanning luminance variations between 0% and 100%. For normalization, this stimulus was divided by the standard deviation of trials across sessions. Conversely, the “stimulus side” parameter characterized the mouse’s behavior upon receiving a reward: continued execution of the same choice after reward receipt or change in behavior in its absence. This binary stimulus side was represented as -1, +1. Furthermore, “past choice” was established as a binary variable, taking values of -1 or 1, denoting left or right prior choices by the mouse, respectively. The “previous reward” (pr) value was defined as -1 or 1, corresponding to incorrect or correct decisions, respectively. Moreover, three “bases” were introduced as linearly independent vectors for the initial 100 trials of each session, capturing the animal’s warm-up effect within the model. In the course of this study, the term “filtered covariate” was employed to reference a covariate subjected to exponential filtering, facilitating the consideration of a temporally filtered variant of the regressor.

Fig. 1d illustrates the distribution of biased data blocks within the IBL experiment, specifically focusing on blocks that extend beyond a defined length ‘n’. These extended biased blocks varied in duration, spanning from 20 to 100 trials, with an average length of approximately 58 trials. It’s important to note that a typical experimental session consists of several such data blocks.

The adopted GLM-HMM, described by Fig. 1e, employs a HMM, featuring distinct weight sets for transition and observation components. Therefore, as mentioned, for the examination of the animal’s decision strategies, a Bernoulli GLM is applied to map the animal’s binary decisions to weighted representations of covariates. Also, there is a multinomial GLM to estimate subsequent state probabilities. Notably, the GLM-HMM framework’s strength lies in its allocation of a multinomial GLM to each state, facilitating the intricate depiction of relationships between transition covariates and their associated probabilities.

### The 4-state model best describes animal decision-making strategies

To identify the optimal model, we conducted a thorough comparison of the GLM-HMM with varying numbers of states and employed cross-validation to select the most descriptive model. After evaluating the log-likelihood values, the GLM-HMM with four states demonstrated notably superior performance compared to the 1-state model, making it the best-fitting representation for the experimental data. These four distinct states correspond to four distinct decision-making strategies. The findings suggest that mice consistently employed all states for multiple consecutive trials, with each session seeing the utilization of various states. It was observed that in the GLM-HMM fit of both unbiased and biased data, covariates such as the stimulus (Δ contrast), past choice, stimulus side, and others played crucial roles in predicting the animal’s choice.

To conduct this comparison, following the fitting of the model with different numbers of states, we computed the log-likelihood of the test data. The EM algorithm was employed for the fitting procedure on the training data sessions. Subsequently, in the testing step, the likelihood of the remaining data was calculated based on the model parameters obtained from the training procedure. By summing over all states, the log-likelihood of the test data is stated as:

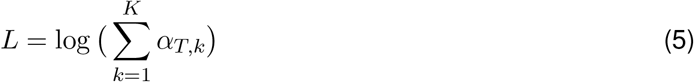

In which *α*_*T,k*_ is the posterior probability of the mice decisions from trial 1 to *T* and was computed solely on the held-out sessions. We can express *α*_*t,k*_ as:

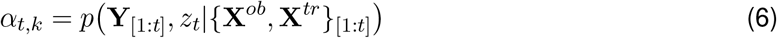

First we performed the global fit in which, the model was applied globally to the entire IBL dataset, and the normalized log-likelihood (NLL) for this data was calculated. The defined normalized log-likelihood, measured in bits per trial (bpt), provides a more intuitive understanding of the model’s performance and facilitates meaningful comparisons across various models and dataset. This can be expressed as:

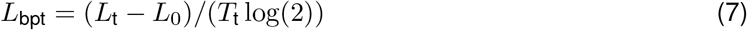

In this equation, *L* is defined as above, *L*_0_ represents the log-likelihood of the baseline model, and *T*_t_ is the number of trials in the test set. The equation has been divided by *T*_t_ log(2) to present the value per trial. The difference between *L* and *L*_0_ represents the enhancement in log-likelihood due to the performance of the GLM-HMM.

Analyzing the Test NLL plot (Fig. 2a), it is evident that the four-state model demonstrates better performance and interpretation, particularly when considering the structured nature of both biased and unbiased data. So, derived GLM weights within the context of the four-state model unveil distinct patterns which are shown in Fig. 2.

**Figure 2:**
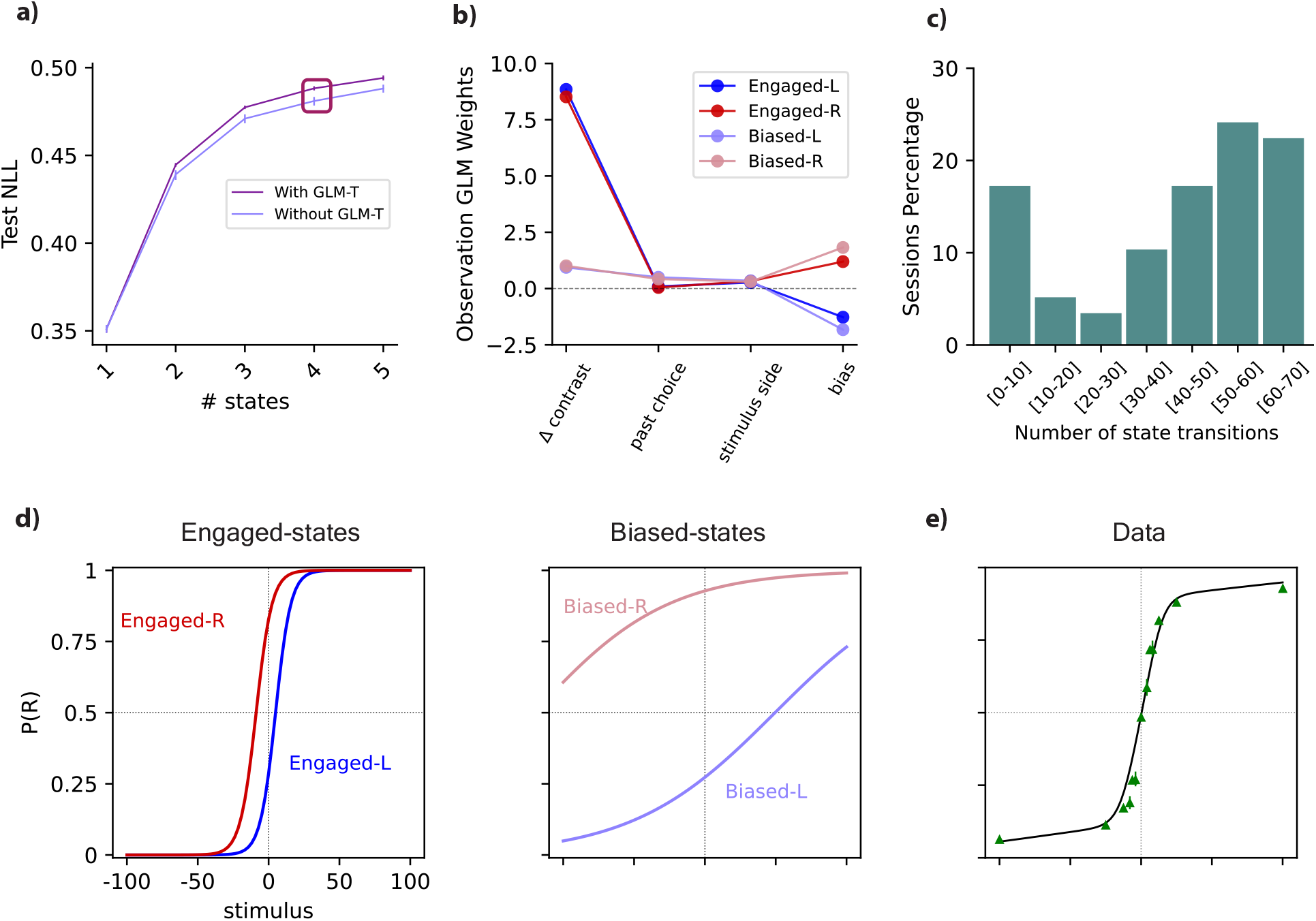
Analysis of the GLM-HMM framework applied to the pooled IBL data. **(a)** A comparison between the Test normalized log-likelihood (NLL) of the model without transition GLM (multinomial GLM) and the model incorporating both GLM-O and GLM-T. The NLL exhibited improved performance in our model with the inclusion of GLM-T, in contrast to the model lacking a transition GLM. **(b)** The inferred observation weights within the four-state model unveil distinct patterns. State 1 demonstrates a substantial stimulus (Δ contrast) weight alongside a moderate left bias, representing the “Engaged-L” state. Conversely, state 2, the “Engaged-R” state, showcases significant stimulus and moderate right bias weights. In state 3, “Biased-L”, and state 4, “Biased-R”, stimulus weight diminishes while bias weights induce strong leftward and rightward biases respectively. **(c)** The histogram illustrates the distribution of inferred state changes per session across sessions exceeding a duration of *T*. Here *T* is the average length of the animal’s sessions, which was equal to 821 trials. **(d)** State-specific psychometric curves for a four-state GLM-HMM. **(e)** The standard psychometric curve derived from the entirety of choice data attributed to this mouse emerges as a composite manifestation, originating from the amalgamation of the four per-state curves depicted in d. The green triangles represent the mouse’s experimental choice data.

In Fig. 2a, for comparative analysis of different models, we computed the log-likelihood for all animal data. The computation strategy involved 5-fold cross-validation with reserved sessions. In this model, when evaluated on the held-out data, the four-state GLM-HMM performed much better than the GLM itself (1-state model). Additionally, the GLM-HMM successfully captured the temporal pattern of inhibition influencing the animal’s decision-making process. Also, a comparison was drawn in terms of the Test Normalized Log Likelihood between the model solely incorporating the Bernoulli GLM and the model encompassing both the GLM-O (Observation) and GLM-T (Transition). This is presented in Fig. 2a.

Our analysis revealed an enhanced performance of normalized log-likelihood (NLL) in the presence of the transition GLM (GLM-T) within our model, as opposed to the scenario wherein the transition GLM was absent. Notably, in our model (purple color, labeled by “with GLM-T”), the 4-state model yielded a 0.01 bps (bits per session) improvement in log-likelihood in comparison with the model without GLM-T. This shows that by considering a dataset with 1000 trials, it is approximately 1024 times more probable that this data is generated by the GLM-HMM with GLM-T than the model without a GLM for transition.

In Fig. 2b, state 1 exhibits a substantial weight attributed to the Δ contrast (stimulus) and a moderate weight associated with left bias, characterizing it as the “Engaged-L” state. State 2, designated as the “Engaged-R” state, features a significant weight related to the stimulus and a moderate right bias weight. Conversely, states 3 and 4 display diminished stimulus weights. However, bias weights induce strong left bias for state 3 and right bias for state 4, denoted as “Biased-L” and “Biased-R” respectively. In these latter states, a nominal weight is also assigned to the previous choice factor.

Also, a histogram was constructed to depict the frequency distribution of inferred state changes per session across all 58 sessions of data for this mouse in Fig. 4c. In Fig. 4c, we specifically analyzed sessions encompassing the initial *T* trials within sessions with lengths more than *T* trials. In this context, *T* denotes the mean duration of all sessions conducted for the respective animal and it was 821 trials. It’s noteworthy that in approximately 18% of all sessions, the mouse exhibited fewer than 10 state changes, resulting in a state duration lasting for an average of 154 trials (calculated by dividing 821 trials by 5 state changes). This observation underscores a consistent tendency for this mouse to remain in specific states for extended periods, particularly the high-performance states. These extended periods of stability were often punctuated by state changes, which primarily occurred when the mouse either adopted a new strategy or shifted its attention to different covariate effects.

On the other hand, in the majority of sessions, approximately 63%, the mouse underwent more dynamic behavior with more than 40 state changes, averaging around 15 trials per state (derived by dividing 821 trials by 55 state changes). This extensive state-switching behavior reveals a decision-making process, where the mouse continuously adapted its strategies and attention throughout the session. This substantial variation within and between sessions dispels any notion that these states merely represent different strategies employed on separate days or sessions.

The psychometric curve, a fundamental tool in psychophysics and decision-making modeling, is typically characterized by a sigmoid-shaped function. This sigmoid-shaped function is intricately linked to a linear representation of the stimulus (Δ contrast) and augmented by a bias term. This mathematical framework is widely adopted to capture the relationship between stimuli and an individual’s responses. Here, the psychometric curve graphically represents the choice probability (right side) relative to stimulus contrast. State-specific psychometric curves were meticulously generated within the framework of a four-state GLM-HMM presented in Fig. 2d. The resulting curves serve as intricate depictions of the behavioral responses observed within each state. The psychometric curves for Biased-L and Biased-R states exhibit shallower inclines, indicative of notable leftward and rightward biases, respectively.

In Fig. 2e, the green triangles correspond to the experimental choice data of the mouse (alongside 95% confidence intervals). Also, to derive the solid black line, a temporally sequenced dataset was generated to match the trial count of the example mouse. This process involved using the meticulously fitted parameters of the GLM-HMM specific to this animal and the actual sequence of stimuli presented during its trials. For each trial iteration, the probability (p(right)) was calculated for each of the nine potential stimuli, regardless of the actual stimulus presented. This calculation entailed averaging the per-state psychometric curves, as illustrated in Fig. 2d while adjusting their weights according to the pertinent row in the transition matrix as this adjustment was contingent on the latent state sampled in the preceding trial.

### Exploring the dynamic transitions and temporal behaviors within the model

Fig. 3a represents both the stimulus and the animal’s choice on the same trial, complemented by the transition regressors of the model. These transition regressors include filtered stimulus side, filtered choice, and filtered reward, providing a comprehensive illustration of the multinomial GLM inputs in the model.

**Figure 3:**
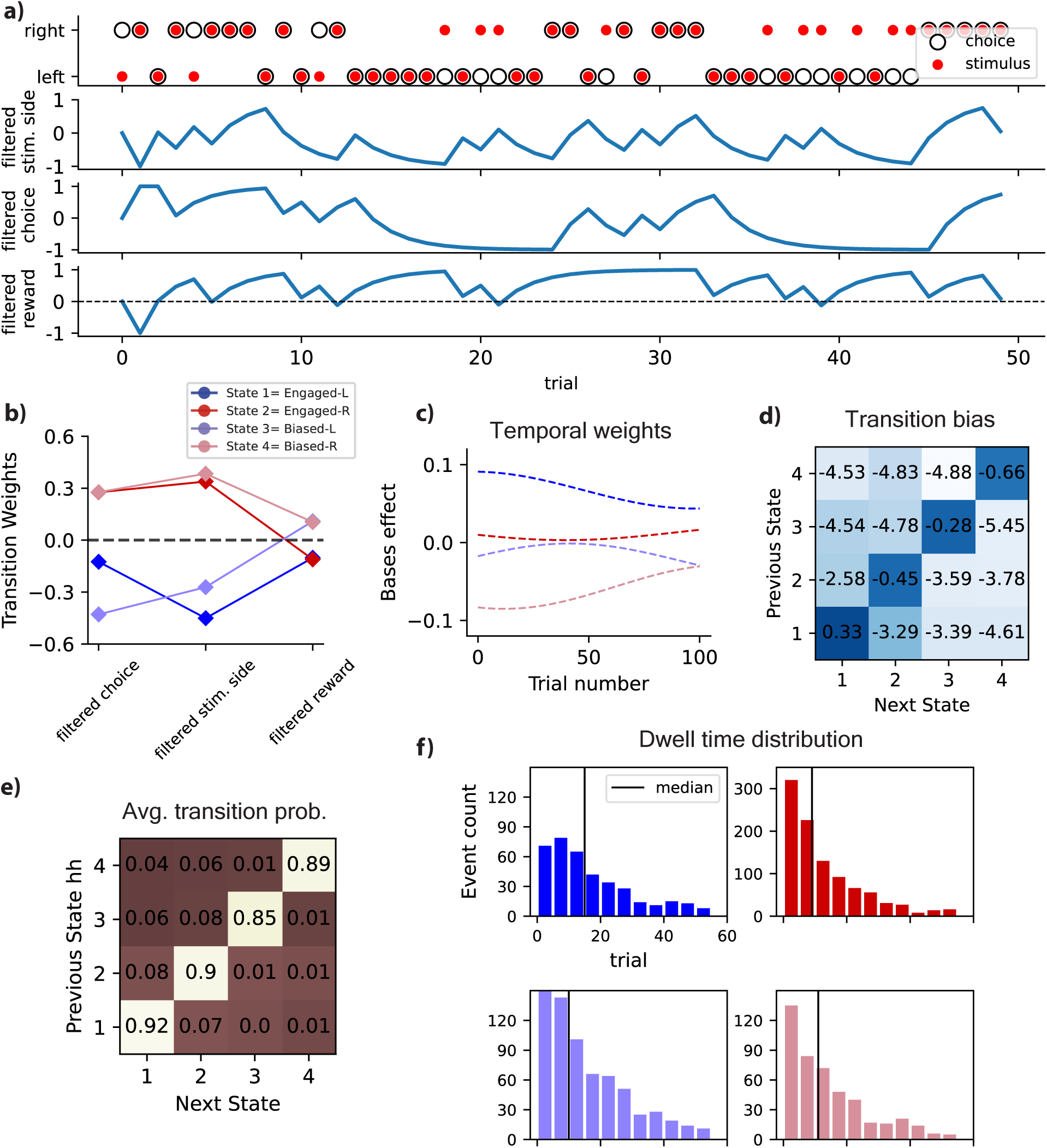
Transitional and temporal dynamics of the model. **(a)** The stimulus (Δ contrast) and animal’s choice, along with the transition regressors of the model, encompassing filtered stimulus side, filtered choice, and filtered reward. **(b)** The transition weights associated with the multinomial GLM. Negative weights for the filtered choice and filtered stimulus side in states 1 and 3 point to left-biased transitions, while positive weights in right-biased states (2 and 4) suggest a preference for right-oriented choices. Disengaged states (3 and 4) exhibit a positive previous reward weight, contrasting with engaged states (1 and 2), where the stimulus holds sway, resulting in a negative previous reward weight. **(c)** Temporal modulation of the bases effect which results from the multiplication of bases weights by bases traces (Three “bases” were introduced into the model to capture the animal’s warm-up effect during the first 100 trials of each session.) **(d)** The transition bias of the model for pooled IBL data. **(e)** The deduced transition matrix for the optimal four-state GLM-HMM, tailored to the entirety of mice IBL data. This matrix displays prominent values along the diagonal, indicative of a pronounced likelihood of persisting within the same state. **(f)** The anticipated duration of stays in different states for all mice: this was achieved by utilizing the derived transition matrix for each individual mouse. Dwell Time Histograms for Engaged-L, Engaged-R, Biased-L, and Biased-R states respectively

The multinomial GLM weights for transition between states are shown in Fig. 3b. For states 1 and 3, characterized by a left-biased component, negative weights are assigned to the previous choice and previous stimulus side. This suggests that these factors contribute to the transition towards left-biased states (Engaged-L or Biased-L). Conversely, for the right-biased states, including states 2 and 4, positive weights are assigned to the previous choice and previous stimulus side, indicating a tendency towards right-oriented choices. Additionally, in disengaged states (states 3 and 4), the positive weight of the previous reward signifies its role in decision-making, as the animal is less attentive to the stimulus in these states. This pattern is reversed in engaged states (states 1 and 2), where the stimulus is pivotal, resulting in a negative weight for the previous reward. Notably, the term “filtered covariate” here refers to transition regressors subjected to exponential filtering, facilitating the integration of temporally filtered versions of these regressors in the analysis.

Also, a collection of three “bases” was introduced, each comprising linearly independent vectors. Temporal modulation of the bases effect is presented in Fig. 3c. The bases were strategically integrated to encapsulate the gradual adaptation of the animal during the initial 100 trials of every session, effectively capturing the warm-up effect within the model.

Fig. 3e presents the inferred transition matrix pertaining to the most suitable four-state GLM-HMM, designed to accommodate the comprehensive mice IBL dataset. Evident within this matrix are notable magnitudes along the diagonal, serving as clear indications of a heightened probability for the system to sustain its presence within the same state during transitions. This prominence underscores the significant propensity for the model to exhibit persistence and stability within individual states.

Furthermore, our analysis delved deeper into the dynamics of state transitions by utilizing the diagonal components within each transition matrix. These components allowed us to calculate the expected duration of residence, referred to as dwell time, for each animal within distinct states, as illustrated in Fig. 3f. The results of this analysis revealed intriguing insights into the temporal aspects of decision-making behavior. Specifically, the median duration of residence in the Engaged-L and Engaged-R states was approximately 10 trials, signifying that the animal spent a reasonable number of trials in these states. Moreover, for the Biased-L and Biased-R states, the median dwell time was found to be 15 and 12 trials, respectively. This observation indicates that the animal exhibited extended periods of engagement in all four states, with slightly longer duration noted in the biased states. These insights shed light on the persistence of specific behavioral states and provide valuable information about the temporal dynamics underlying decision-making behavior in mice.

### Apply the model for the interpretation of a mouse data example

The findings for a specific mouse of IBL data as a case study are presented in Figures 3 and 4, providing a detailed insight into an individual fit. To delve into the temporal aspects of decision-making behavior, we harnessed the power of the GLM-HMM we fitted. This model allowed us to calculate the posterior probability of the mouse’s hidden state throughout all the trials. These state trajectories essentially represent our best estimates of what the mouse’s internal state likely was on each trial, considering the entire sequence of observed inputs and choices during a session. The plots showcase the posterior probabilities and correct/incorrect animal choices associated with their corresponding sessions in four distinct states, as presented in Fig. 4a. These visualizations incorporate a background color scheme that serves to distinguish biased blocks: the color pink corresponds to right-biased blocks, while the blue shade denotes left-biased blocks. This color-coded representation enhances the comprehensibility of the data, facilitating the identification of patterns within the biased segments of the experiment. These graphical representations illustrate that states exhibiting right bias, namely Engaged-R and Biased-R, demonstrated greater likelihood within blocks characterized by a right-biased orientation (designated by a light pink background color). A similar observation was made for left-oriented blocks with a blue background hue, where states influenced by left bias exhibited higher probabilities. Also, as we can see Fig. 4a, what emerged is a clear pattern: one state consistently stood out as significantly more probable than the others, signifying a strong level of confidence in our understanding of the mouse’s internal state based on the observed data.

**Figure 4:**
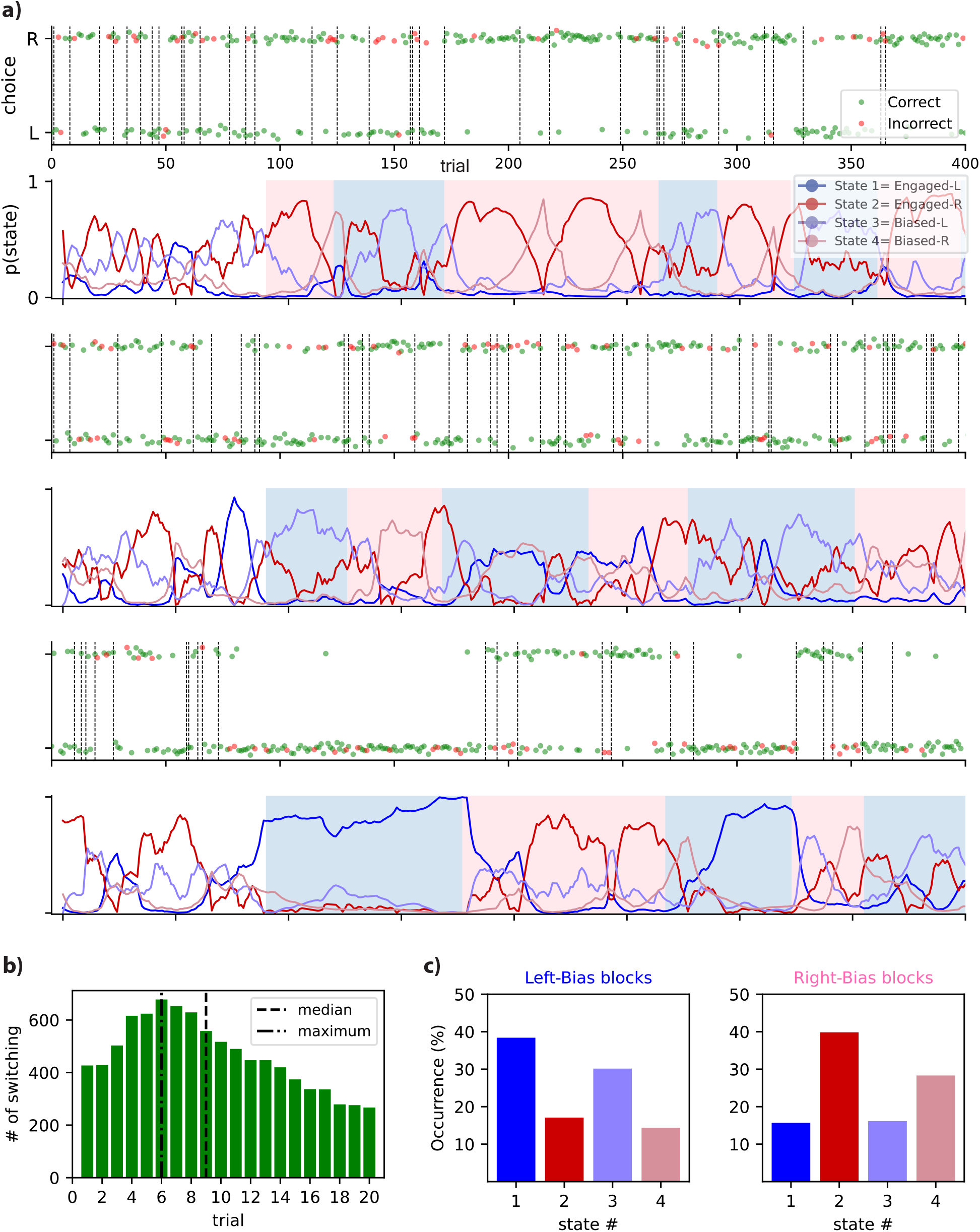
Interpretation of mouse data using GLM-HMM. **(a)** The plots display posterior probabilities and correct/incorrect decisions of mice while performing the task for four distinct states. The background color distinguishes biased blocks: pink indicates right-biased blocks, while the blue shade represents left-biased blocks. **(b)** The histogram depicting the frequency of initial transitions into corresponding states were generated for each block (e.g. initial transition to states with a right bias for right-biased blocks of data). **(c)** An evaluation of the fractional occupancy pertaining to the four discrete states. This was conducted over the entire span of trials encompassing both right-biased and left-biased blocks. The results are presented here separately for the two different data block types.

A histogram was meticulously constructed to elucidate the distribution of the first transition into states aligning with related bias within each data block (Fig. 4b). This analytical approach is geared towards capturing the number of trials spanning from the commencement of a biased data block until the occurrence of the first transition to the corresponding biased states. For instance, in the context of a right-biased data block, this entails quantifying the number of trials required for transitions to the states characterized by a right bias, namely Engaged-R, and Biased-R. A parallel assessment is made for left-biased data blocks and their respective states with a high value on the left bias weight. The analysis underscores that the median value of this histogram stands at 9 trials, while the maximum value observed is 6 trials. This implies that, on average, it takes approximately 9 trials for mice to transition from the start of a biased data block to a state matched with the prevailing bias.

Consequently, this model provides a profoundly distinct perspective on mouse decision-making behavior. It not only captures the temporal dynamics but also reveals the diverse patterns and strategies that mice employ during decision-making processes, offering a more comprehensive and nuanced understanding of their cognitive processes.

In Fig. 4c we conducted an assessment of the fractional occupancy concerning the four distinct states across the entirety of trials for right-biased and left-biased data blocks. In this analytical endeavor, we assigned each trial to the state that exhibited the highest likelihood and proceeded to compute the proportion of trials designated to each respective state.

As evident from the analysis, for each bias-related plot, the corresponding states were observed to encompass a substantial portion of the entire trial set, indicating a significant representation of the mouse’s behavioral responses within biased data blocks. In the context of right-biased data, the right-side plot in Fig. 4c, Engaged-R and Biased-R (represented by red and pink columns) exhibited notably higher values in comparison to the other associated columns. A similar observation was made for left-biased blocks (left-side plot in Fig. 4c), where Engaged-L and Biased-L showed elevated values compared to the other two states. This observation underscores the impact of bias weights on biased data blocks, highlighting the adaptive nature of the mouse’s decision-making process within the experimental framework.

Remarkably, the analysis uncovered that the mice spend approximately 72% of their time in related biased states (e.g., Engaged-R and Biased-R in right-biased blocks), with the engaged state having a slightly higher occurrence chance. As mentioned, this intriguing revelation was derived from an extensive dataset comprising 37 mice. In stark contrast, the mice allocated a relatively smaller fraction of their trials, approximately 28%, to the other unrelated states (e.g., Engaged-R and Biased-R in left-biased blocks).

### Analyzing each individual animal yields consistent findings

To gauge the universality of these discoveries, we applied the GLM-HMM model to the choice data obtained from all animals in the IBL dataset (37 individual fits for all mice). Therefore, two sets of GLM weights, observation, and transition weights, for these animals are presented in the Fig. 5 a,b. A notable level of substantial agreement was distinctly apparent upon analyzing the fits of the four-state GLM-HMM in the study. This consensus was particularly pronounced, as a significant majority of the mice showcased discernible states, identified as ‘Engaged-L’, ‘Engaged-R’, ‘Biased-L’, and ‘Biased-R’ (Fig. 5 a,b). This alignment in the characterization of states underscores the robustness and consistency of the applied GLM-HMM framework in capturing these behavioral patterns across the population of interest for both observation (Bernoulli GLM) and transition weights (multinomial GLM).

**Figure 5:**
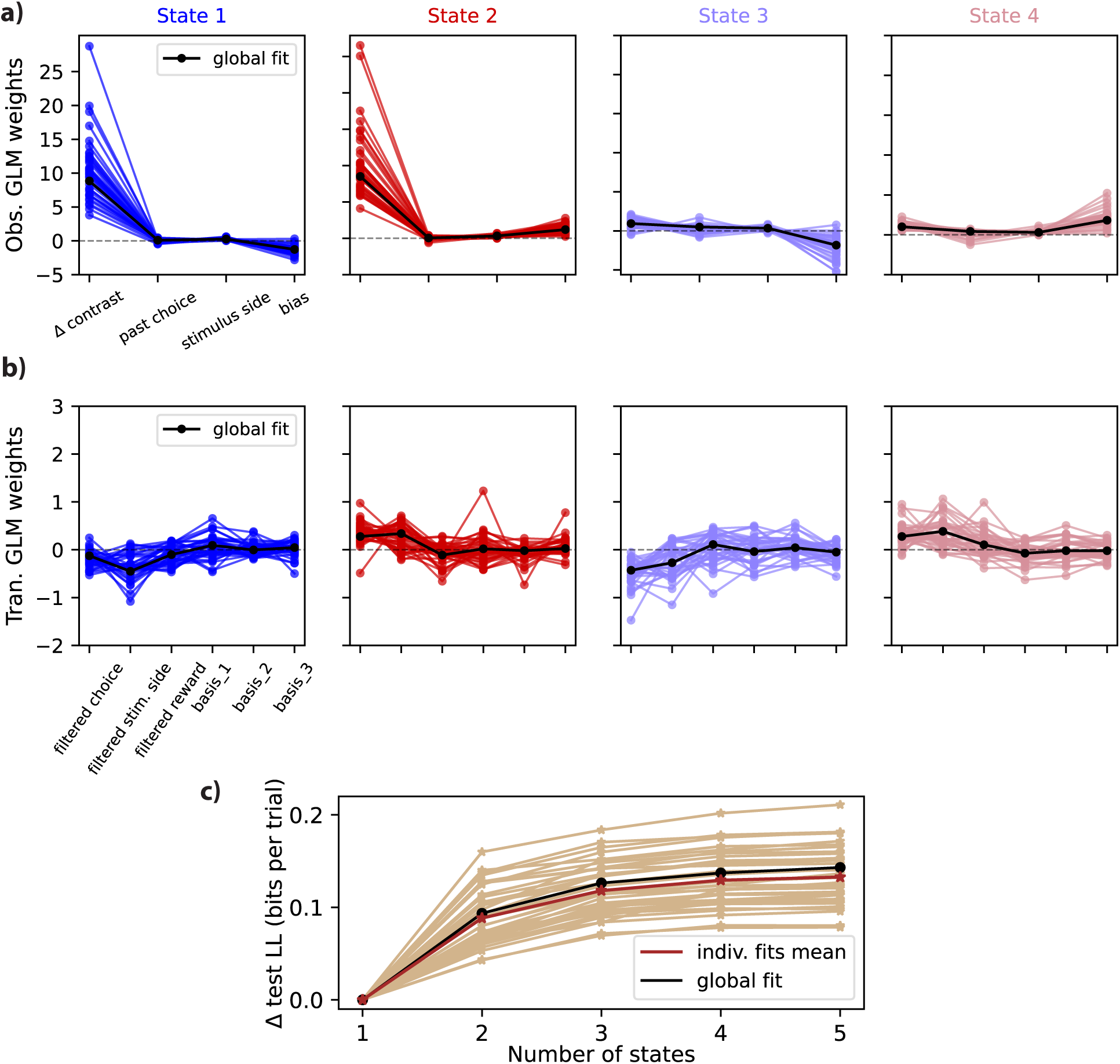
Analysis of data from each individual mouse in the IBL dataset. **(a)** Observation weights for Bernoulli GLM corresponding to distinct states within the four-state model across all IBL animals. **(b)** multinomial GLM transition weights pertaining to diverse states within the four-state model across all IBL animals. **(c)** For each mouse in the population, the test log-likelihood variation relative to a (one-state) GLM is illustrated against the number of states. Each individual mouse is represented by a distinct trace.

The Fig. 5c presents an analysis of the test log-likelihood variation in relation to the number of states, specifically pertaining to each mouse within the examined population. Each distinct trace on the graph represents an individual mouse’s data, offering a holistic view of the log-likelihood changes across different state configurations. The solid black lines effectively visualize the collective mean across all animals, providing a clear reference point for comparison. The findings presented in Fig. 5c align with the trends observed across the entire cohort of animals, suggesting a robust pattern. To provide further detail, it’s noteworthy that the four-state GLM-HMM consistently displayed superior performance when compared to the single-state GLM during cross-validation. This trend held true across the entire group of 37 mice included in our study. This consistency in results strengthens the reliability and validity of our multi-state GLM-HMM framework with two sets of GLM as a powerful tool for understanding decision-making behavior in mice across diverse individuals.

## 3 Discussion and conclusion

In summary, this paper presents a GLM-HMM framework for modeling and analyzing mice’s decision-making behavior in non-stationary environments. This model offers a more nuanced understanding of transitional patterns of behavioral states. This model is structured as an HMM, featuring two sets of per-state GLMs, referred to as GLM-O for observations and GLM-T for transitions. This design provides flexibility in capturing the impact of covariates on both mouse choices and state transitions. When applied to the IBL dataset, our analysis demonstrates the superior performance and interpretability of the four-state GLM-HMM compared to the 1-state model. This four-state model effectively uncovers intricate patterns within the data, with states characterized by bias weights adeptly capturing changes in biased data blocks. The transition and observation GLMs contribute significantly to representing the animal’s choices and transition patterns accurately. In essence, our study underscores the GLM-HMM’s effectiveness in modeling animal decision behavior and transition probabilities between states, especially in scenarios with variable stimulus probabilities.

While our discussions have primarily focused on the specific 4-state model, it’s crucial to emphasize that the conclusions and findings presented here are not limited to this configuration alone. Our results and insights can be generalized to a 5-state model (in Supp.). Although the cross-validated log-likelihood gain is slightly higher for the 5-state model, the results obtained remain similar in several aspects.

Looking ahead, it would be useful to have a look at whether there are any clusters that can be seen in the individual fits for different animals. For example, are some mice less engaged than others, and do these engagement rates persist across sessions? Furthermore, it would be valuable for future research to explore a comparative analysis between the discrete state model presented in this paper, the GLM-HMM, and a model that incorporates continuously changing states over time. This comparison could shed further light on the intricacies of decision-making behavior in animals and deepen our understanding of the underlying neural processes.

Also, the future holds exciting prospects for advancing our understanding of neural activity in relation to the detected discrete behavioral states. Unraveling the intricate patterns of neural and behavioral correlations promises to be a pivotal direction in neuroscience research. By harnessing advanced techniques, such as functional magnetic resonance imaging (fMRI) and electrophysiological recordings, researchers can delve deeper into the neural underpinnings of animal decision-making behavior. Identifying specific neural signatures associated with each state could provide unprecedented insights into the mechanisms driving these behavioral strategies. This endeavor may uncover patterns of neural activity that precede or coincide with state transitions, shedding light on the neural mechanisms that drive shifts in decision strategies.

On the other hand, it’s worth noting the broader landscape of hierarchical models for analyzing behavioral data and decision-making in mice. Hierarchical models offer a powerful mean to capture the complexity of behavioral data by incorporating multiple levels of variability and structure. In the context of decision-making, these models can account for individual differences in behavior, learning, and strategy adoption. Furthermore, they allow for the integration of neurophysiological data, enabling a deeper understanding of the neural underpinnings of decision-making processes. Exploring the synergy between hierarchical modeling and GLM-HMM techniques holds promise for unraveling the intricate relationships between neural activity, discrete behavioral states, and the decision-making strategies employed by animals.

## 4 Methods

### 4.1 Hidden Markov Model (HMM)

A Hidden Markov Model (HMM) is a statistical model for time series data that is governed by hidden or “latent” factors that cannot be directly observed. The events that we observe are called observations, and the underlying, unobservable factors driving them are referred to as latent states [18–21].

Therefore, HMM consists of two stochastic components: one governing the latent states, and another governing the observations. The latent process component satisfies the Markovian property. A transition probability matrix *A ∈* ℝ^*K×K*^, where, at trial *t*, the element corresponding to state *j* and state *k* presents the transition probability between those two states and can be written as:

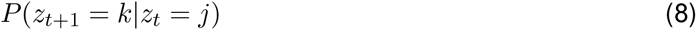

and an observable state-dependent component 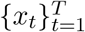, where

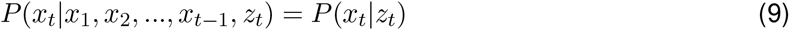

where *T* is the number of considered trials. Generally, in HMM, the probability distribution of the observed symbols is based on the underlying, unobserved states of the system, following the principles of a Markov chain. In many cases, observations can be grouped into different classes, which can provide more insightful information than the individual observations themselves. In such situations, it becomes advantageous to model these observations using both the observable and unobservable aspects of HMM.

### 4.2 Bernoulli GLM

The Bernoulli Generalized Linear Model (GLM) is a statistical model designed for binary data, where the response variable can take values of 0 or 1. It belongs to the broader GLM family, which includes models like Poisson, Gaussian, and Dirichlet. The primary purpose of using Bernoulli GLMs here is to model the relationship between a mouse’s expected decision and the relevant regressors for each trial.

In the Bernoulli GLM, the response variable follows a Bernoulli distribution, which is a discrete probability distribution. It takes the value 1 with probability *p* (representing success) and 0 with probability 1 *− p* (indicating failure). Estimating the probability *p* in the Bernoulli GLM involves predictor variables, and a link function connects the mean of the Bernoulli distribution to the linear predictor.

The linear predictor is formed as a linear combination of the predictor variables and their respective coefficients. Subsequently, the link function is used to transform the linear predictor to the probability scale. The logit link function stands as the most widely adopted link function for a Bernoulli GLM, and it can be expressed as log(*p/*(1 *− p*)) = *Fβ*, where *F* corresponds to a predictor variable matrix, and *β* represents a vector of coefficients.

In this study, we employed a Bernoulli GLM to analyze the animal’s strategies in relation to various experiment regressors. It can map the binary values of the animal’s decision to the weighted representations of the considered covariates. These weights serve to depict the inputs of the model in relation to the output, which is the animal’s choice on each trial. Consequently, we can describe an observational GLM using the following equation, where the animal choice, denoted by *y*, can take a value of 1 or 0, indicating the mouse turning the wheel to the right-side or left-side, respectively:

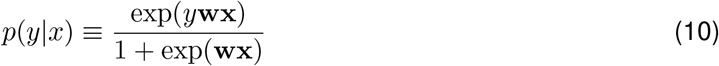

In this equation, as indicated by the notations, the presented GLM is solely associated with the observation covariates, **x**, and observation weights **w**.

The fitting procedure of the model involves utilizing a penalized maximum likelihood estimation. This estimation minimizes the sum of the priors on the transition and observation weights, in addition to a negative log-likelihood function, often referred to as the log-posterior. The prior corresponds to a normal distribution over the weights with a mean of zero and a variance of *σ*^2^. The negative log of this prior can be represented as 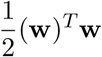. The purpose of this prior is to impose a penalty on the model weights, thereby regularizing the model by discouraging excessively large weight values for the regressors [22]. Consequently, a relevant loss function can be defined as:

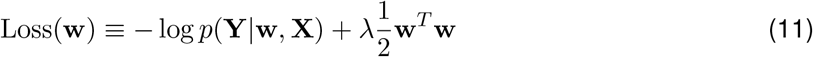

In this equation, log *p*(**Y**|**w, X**) represents the conditional probability of the output, which corresponds to the decisions made by the animals, given the model regressors. Also, the symbol *λ* assumes the role of a hyperparameter that governs the regularization term’s influence on the model. The log-likelihood function can be mathematically defined as follows:

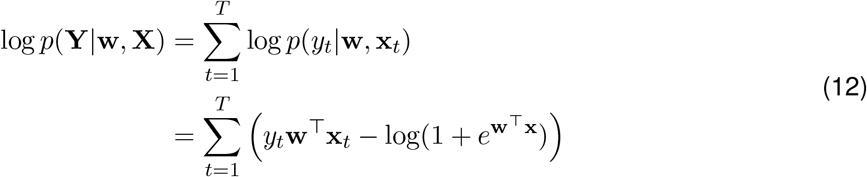

In this context, **Y** represents the observations from trial 1 to *T*, while **X** corresponds to the regressors for the GLM applied to the same trials.

The GLM can be fitted using the maximum likelihood or maximum a posteriori (MAP) estimation, and the resulting coefficients can subsequently be used to make predictions on new data. To assess the performance of the fitted model, we use cross-validation on held-out test data.

### 4.3 multinomial GLM

Another form of the Generalized Linear Model is the multinomial GLM, which we will employ to model how external covariates influence transitions between different states. This multinomial logistic regression, also known as softmax regression or maximum entropy classifier, serves as an extension of logistic regression to handle data with multiple categories. Multinomial GLMs are generalized linear models possessing the capability to analyze data from more than one category simultaneously. By modeling the relationship between independent variables and categorical dependent variables, this approach allows for the determination of the likelihood associated with each category. In the field of neuroscience, multinomial GLMs are frequently employed to analyze the link between brain activity and behavior, facilitating the investigation of neural processes underlying different types of behavior.

The primary difference between Bernoulli and Multinomial GLM lies in the number of categories the models handle. Bernoulli GLM is used for binary outcomes, modeling the probability of success (1) or failure (0), commonly employed in binary classification. Multinomial GLM, on the other hand, deals with data featuring more than two categories, modeling the probabilities of each category using a multinomial logistic function. The choice between them depends on whether you’re working with binary or multiclass categorical data. In multinomial GLM, the expression for the conditional probability of observing a particular outcome **y**, given input variables **x** and a set of model parameters **w**, is typically formulated as follows:

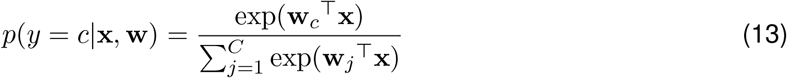

where **w**_**j**_ corresponds to the parameters associated with j-th outcome and *C* is the number of possible outcomes in which for multinomial GLM *C* is more than 2. The multinomial equation finds frequent application within the domain of multinomial logistic regression, serving as a means to model the probability distribution across numerous discrete outcomes or categories. This mathematical framework is foundational in a multitude of machine learning and statistical contexts, including but not limited to tasks such as text classification, image recognition, and addressing multiclass classification challenges.

In this paper, we delve into the exploration of the GLM-HMM, Generalized Linear Model-Hidden Markov Model, with multinomial GLM outputs, a method capable of estimating the likelihood of the next state. Notably, the GLM-HMM framework offers the advantage of enabling each state to possess a multinomial GLM that effectively captures the intricate relationship between transition covariates, such as previous choice and previous reward, and the associated transition probabilities.

Therefore, we consider the animal’s behavior at trial *t* and proceed to calculate transition probabilities to various states at trial *t* + 1 utilizing a multinomial GLM. The model is adeptly structured to establish associations between the vector of model transition inputs and the unnormalized log probability of each potential future state. So our model presents transition probabilities using a multinomial GLM, and emission probabilities using a Bernoulli GLM, each defined by its own set of parameters.

### 4.4 Standard GLM-HMM

GLM-HMM, or Generalized Linear Model-Hidden Markov Model, is a sophisticated probabilistic model in which the core structure combines the principles of HMM and GLM. At its essence, GLM-HMM is defined by a dual-layered structure. The first layer comprises a Hidden Markov Model, a stochastic process with hidden states that transition over time. These hidden states capture latent information about the underlying dynamics of a system. The second layer incorporates Generalized Linear Models, which govern the mapping from inputs to outputs, and this mapping is influenced by the current hidden state of the HMM. This dual-layer structure enables GLM-HMM to effectively model complex, sequential data where the relationship between observed outputs and input variables varies depending on the underlying, unobservable state. This capacity to incorporate state-dependent relationships within a probabilistic framework makes GLM-HMM an appropriate tool for analyzing and understanding temporal data in various domains.

In two previous studies, the GLM-HMM with a Bernoulli GLM for observations was introduced to analyze mouse decision-making behavior [1, 23]. That work used an HMM with Bernoulli GLM, as described in sections 1.1 and 1.2, to elucidate animal behavior. In their model, they considered animal choices as the output of an observation GLM, with parameters involving observation weights and a fixed transition matrix.

### 4.5 GLM-HMM with GLM transitions

In this manuscript, we present a GLM-HMM framework that incorporates both GLM observations and GLM transitions. This model enables us to capture the temporal patterns of animal transitions between states, enhancing our ability to understand the dynamics of decision-making behavior in mice. Therefore we have an additional component here - the multinomial GLM. This component plays a crucial role in capturing the intricate patterns associated with state transitions. So, instead of a fixed transition matrix between states, our model utilizes a set of vectors, one for each state, to capture transitions into that state.

The GLM-HMM with GLM transitions is composed of several key components. These components include the observation model, transition model, emission model, initial state probabilities, and the hidden state sequence. The observation model establishes the relationships between observed and predictor factors. Typically, this component takes the form of a GLM, which models the conditional distribution of the observed parameter based on its predictors. The transition model characterizes the transition probabilities between the various hidden states within the model. At each trial or time point, these probabilities govern the likelihood of transitioning from one state to another.

Therefore, in the context of a multinomial GLM, for the transition model, the GLM output reflects different probabilities for transitions between states. In this case, 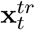 presents the transition covariates at trial *t* and 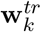 is the transition weights associated with state *k*. So for the transition probability to state *k* at trial *t*, denoted as *z*_*t*_, we can write:

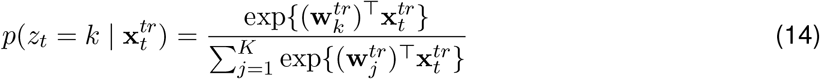

Here, *K* represents the total number of states.

The log-likelihood function in a Multinomial GLM is rooted in the multinomial distribution. It quantifies the probability of observing categorical outcomes based on the predictor variables and model parameters. This log-likelihood function is conventionally formulated as follows:

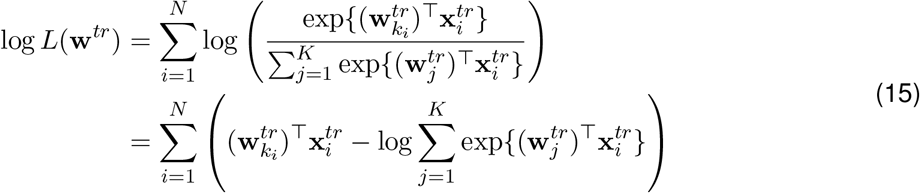

in which *N* is the total number of outputs and 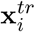 represents the transition regressors for the i-th output. The other details of the mathematical description of this model will be explained in the upcoming sections.

### 4.6 Model inputs

In this framework, we incorporated several covariates into the model, which will be elaborated upon in detail here.

For the GLM observations, the covariates were stimulus, previous choice, previous stimulus and bias. The “stimulus”, in the experiment, was defined as the contrast of a sinusoidal grating, varying between 0% and 100% brightness. To further normalize the stimulus, it was divided by the standard deviation of the trials across sessions. Also, the “past choice” was defined as a binary variable with values of -1 or 1, depending on whether the mouse’s previous choice was left or right, respectively. On the other hand, the “previous stimulus” was defined as the behavior of the mouse when rewarded. If the mouse received a reward, it continued to make the same decision; however, when no reward was given, it changed its behavior. The previous stimulus values were binary, denoted as -1, +1.

The covariates for the GLM transitions included a filtered version of the previous choice, previous stimulus, previous reward, and three basis coefficients. Throughout this paper, the term “filtered covariate” denotes a covariate filtered with an exponential filter, enabling the consideration of a temporally filtered regressor to consider the impact of the regressors in the transition between states. Therefore, the filtered previous choice and previous stimulus are temporally filtered versions of similar GLM observation covariates. The “previous reward” value was set to -1 or 1, based on whether the previous animal’s decision was correct or not, respectively. Furthermore, three “bases” were defined as linearly independent vectors to represent the initial 100 trials of each session. These covariates aimed to capture the animal’s warm-up effect within the model.

### 4.7 Setting a prior on the model parameters

We use Maximum A Posteriori (MAP) estimation to fit model parameters, denoted as Θ = {**w**^*tr*^, **w**^*ob*^, ***π***}. These parameters consist of the initial state distribution, transition weights, and observation weights for all states, which will be explained in more detail in this section.

To implement our approach, we employed the Expectation Maximization (EM) method, as introduced by Dempster et al. [24]. This iterative technique enables the determination of parameters within a given model by maximizing the likelihood of the data, given the specific parameters. Previous works [9, 15, 25] have successfully applied the EM approach to HMMs integrated with external regressors. The parameter estimation using the EM method involves an iterative process with two essential steps: the “expectation step,” where parameter expectations are computed, and the “maximization step,” optimizing data likelihood based on those parameters. This iteration continues until optimal parameter values are determined.

In this study, we incorporated a Dirichlet prior to model the initial state distribution ***π***. As for the GLM, we employed independent zero-mean Gaussian priors for the observation and transition weight vectors, with variances denoted as 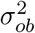 and 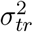, respectively. Larger values of these variances signify a flatter prior distribution.The entire set of model choices and inputs is expressed as *ℱ ≡ {***Y, X**^**ob**^, **X**^**tr**^}. Here, 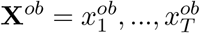 represents the observation weight vectors, and 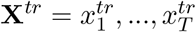 corresponds to the transition weight vectors. The animal choices for the specified trials are denoted as **Y** = *y*_1_, …, *y*_*T*_. The prior distribution considered for our GLM-HMM, *p*(Θ), is as follows:

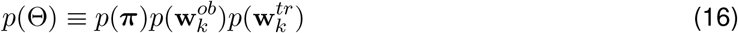

In this context, the model parameters encompass three distinct categories, namely, the initial state distribution, the state-specific observation weights and state-specific transition weights denoted by 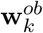 and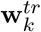, respectively. Here, *k* denotes the number of states, ranging from 1 to *K*. The initial state distribution is represented by ***π*** *∈* ℝ^*K*^. The prior distributions for the observation and transition weights in the GLM are both assumed to follow normal distributions, as expressed in the following equation:

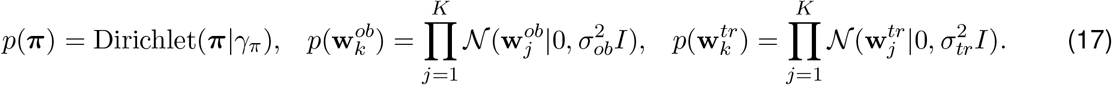

The prior value for the initial state distribution was assigned as *γ*_*π*_ = 1. To determine the optimal hyperparameter value for the model prior on weights, we conducted a grid search over a range of values, specifically *σ ∈ {*0.25, 0.5, 1, 2, 4, 8, 16}. We employed a held-out validation set to compare the Normalized Log-Likelihood (NLL) values for the different *σ* values within the specified set. Subsequently, the value of *σ* yielding the highest NLL on the IBL data was selected, and it was found to be *σ* = 4.

The GLM-HMM employs a hidden Markov model, encompassing distinct sets of weights for both transition and observation aspects. Within the model, each state is governed by a state-specific Bernoulli GLM and a multinomial GLM, representing the animal’s decision-making behavior concerning choice probability and the probability of transitioning between states, respectively. In this context, we used the notations GLM-O and GLM-T to represent GLM observations and GLM transitions, respectively.

The transition multinomial GLM involves weights that associate relevant regressors, denoted as **x**^tr^, with the probabilities of transitioning between states. These transition probabilities are not fixed values and are contingent upon the combination of related regressors and the current state. They are represented by a matrix *A ∈* ℝ^*K×K*^, where, at trial *t*, the element corresponding to state *j* and state *k* presents the transition probability between those two states and can be written as:

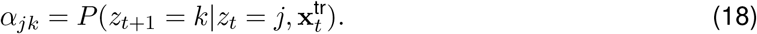

It is pertinent to acknowledge that the observation Bernoulli GLM plays a crucial role in determining the observation weights and characterizing the decision behavior as a function of the input observation regressors.

Ultimately, in the context of mice decisions, along with transition and observation covariates, the primary goal of the EM algorithm is to optimize the log-posterior of the model parameters. The log-posterior can be mathematically expressed as follows:

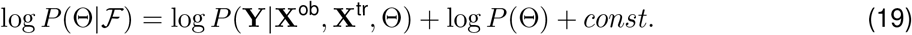

### 4.8 Model Fitting

The GLM-HMM employs the EM algorithm [25] to optimize model parameters for maximizing the likelihood of observed data [22]. This iterative algorithm consists of two main steps: the E-step, where the expected complete-data log-likelihood is computed based on parameter estimates and observed data, and the M-step, which maximizes model parameters based on these expectations. The likelihood is calculated using the forward-backward approach [22], a dynamic programming algorithm. The EM algorithm continues iteratively until convergence is achieved, ensuring an accurate parameter estimation process.

To elaborate further, during each trial and based on the specified GLM-HMM parameters, we compute the joint probability distribution encompassing both the states and the animals’ decisions (left or right). Subsequently, the log-likelihood of the model is evaluated using this joint probability distribution. This relationship can be expressed in the following manner:

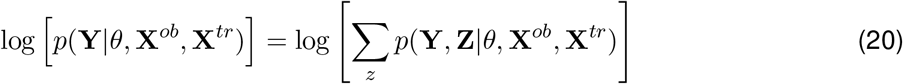

In the context of our model, we define two sets of covariates: 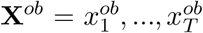 represents the observation covariates, and 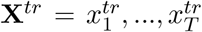 represents the transition covariates. Additionally, we have a set of latent states denoted as **Z** = *z*_1_, …, *z*_*T*_, and corresponding observations for these states denoted as **Y** = *y*_1_, …, *y*_*T*_ .

In the model, **X**^*ob*^ and **X**^*tr*^ capture relevant information related to the observations and transitions, respectively. These covariates play a crucial role in characterizing the underlying dynamics of the system under consideration. The latent states, **Z**, represent unobservable or hidden variables that drive the observed data. They are essential components of the model, as they provide insights into the underlying processes governing the observed phenomena. The observations, **Y**, are the data collected from the system, corresponding to each specific latent state in **Z**. These observed data points are used in the model to estimate and infer the hidden states and the model parameters.

In the GLM-HMM, state-dependent GLM-T weights represents transition regressor weights, and GLM-O weights represents the significance of observation covariates. These weights’ patterns differ across distinct states. In the Bayesian context, where prior information exists for unknown parameters, the EM algorithm can be used to compute the mode of the posterior probability distribution, facilitating parameter estimation.

In the context of probabilistic graphical models and variational inference, the E-step is often associated with variational lower bounds. Specifically, when dealing with intractable posterior distributions or complex models, the E-step aims to maximize a lower bound on the log-likelihood, rather than the log-likelihood itself. This lower bound is often referred to as the Evidence Lower Bound (ELBO) or the variational lower bound. Here, during the E-step of the EM algorithm, a lower bound on the right-hand side of the objective function referred to as Eq. 20 [24, 26] is maximized. Subsequently, in the M-step, the expected log-likelihood obtained from the E-step is maximized with respect to the GLM-HMM parameters. In the following sections, we will detail the calculations for both the E-step and M-step of this estimation approach. These steps play a crucial role in iteratively refining the parameter estimates until convergence is achieved, enabling the determination of the mode of the posterior probability in the Bayesian setting.

By defining the Bernoulli GLM distribution as 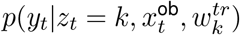, we obtain the following equation for the expected log-likelihood during the E-step:

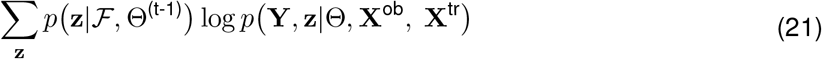

In this equation, we can express the model joint distribution, *p*(**Y, z**|Θ, **X**^ob^, **X**^tr^), as:

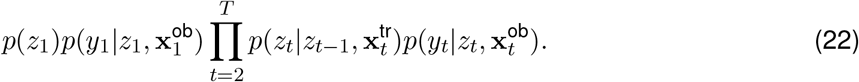

To calculate this expected log-likelihood, the E-step uses a forward-backward approach [14] to estimate the single and joint posterior state probabilities. We are going to explain each one separately here. So the process involves calculating the single posterior state probability at trial *t*, given by:

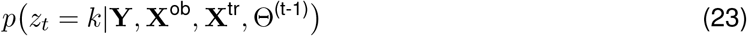

We show this probability by *ϕ*_*t,k*_. In the forward-backward approach, the E-step iteratively calculates the posterior probability of the mice decisions by going through a loop. As a result, the single posterior state probability can be decomposed into the following equation:

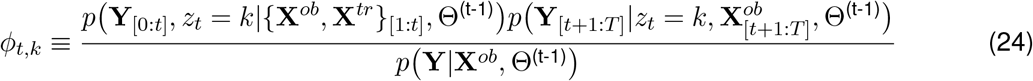

In which Θ^(t-1)^ is the parameters of the model at trial *t −* 1.

In the E-step of the estimation process, the forward-backward algorithm is used to compute the expectation of the desired function. This two-stage message-passing algorithm is also known as the filtering process during the forward pass [25]. The forward-backward algorithm, through its filtering process, computes the posterior probabilities for each time point, enabling the estimation of the latent states’ influence on the observed data. In the following, we will explain the forward and backward passes of the algorithm

#### Forward pass

In the E-step, the forward–backward algorithm is employed to obtain the expectation of the desired function. This is a two-stage message-passing algorithm [25] and the forward pass is called filtering. By assuming we have GLM-HMM parameters and observation data, to calculate the E-step, we should form the posterior probability as a function of latent states.

The objective of E-step is to infer the hidden state sequence, given the available observations and the current parameter estimates. In above equation for the first trial, the posterior probability is obtained by 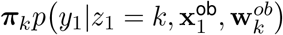.

Here, 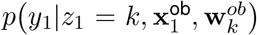 is the Bernoulli GLM distribution for observations (GLM-O). In this equation, if we consider the posterior probability of the mice decisions, *α*_*t,k*_, from trial 1 to *t* as:

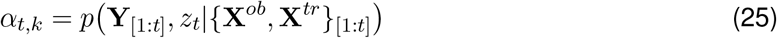

Based on this definition, for the first trial, we can call the posterior probability as *α*_1,*k*_. Then using earlier calculations, for upcoming trials until *T*, we can compute the posterior probabilities, *α*_*t,k*_, as:

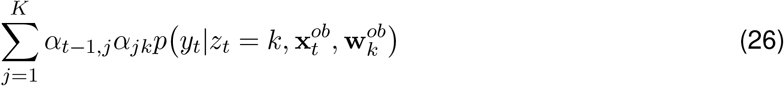

#### Backward pass

In the forward-backward algorithm, the backward step complements the forward pass, utilizing previously obtained information to update latent state probabilities. Named “backward” due to its reverse chronological order, it calculates updates essential for the M-step. The backward pass relies on the joint posterior distribution of two latent states, contributing to refined GLM-HMM parameter estimates. Its primary goal is to determine the posterior probability of observed mouse data for all states and trials beyond the current trial. This probability is represented as *β*_*t,k*_, where *t* denotes the trial index and *k* refers to a specific state. This can be written as:

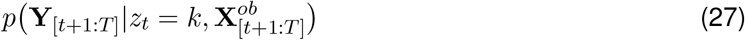

Therefore, for trial *T*, we have *β*_*T,k*_ = 1. Also, we can compute the posterior probability, *β*_*t,j*_, for trials *{T −* 1, …, 1} as:

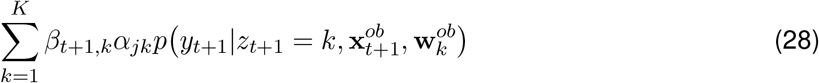

Here, it should be considered that a Bernoulli GLM distribution for observations can be written as 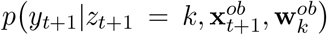. By incorporating the information from both the forward and backward passes, the model gains a better understanding of the system’s underlying dynamics and is better equipped to estimate the latent states’ influence on the observed data, leading to improved parameter estimation in the M-step of the EM algorithm. So, by employing the forward-backward approach, we have:

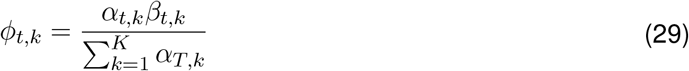

On the other hand, for the joint posterior state probability *μ*_*t,j,k*_, we can write:

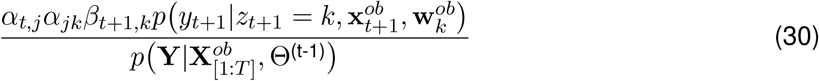

and by substituting the denominator using the definition of posterior probabilities, we have:

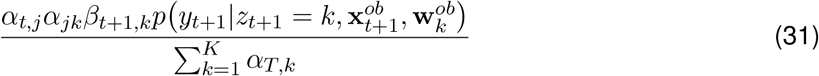

Therefore, to calculate expected log-likelihood, we should use the results of the posterior state probabilities which was obtained using the forward-backward algorithm. Therefore, for a given trial *t* and state *k*, the posterior state probability is defined by *ϕ*_*t,k*_ *≡ p* (*z*_*t*_ = *k*|*ℱ*, Θ^(t-1)^) and the joint posterior state distribution for states at trials *t* and *t* + 1 is given by *μ*_*t,j,k*_ *≡ p* (*z*_*t*+1_ = *k, z*_*t*_ = *j*|*ℱ*, Θ^(t-1)^). In conclusion, by considering the single and joint posterior state probabilities, *ϕ*_*t,k*_, *μ*_*t,j,k*_, we can rewrite the Eq. 21 as:

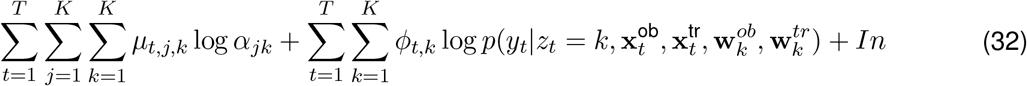

Here *In* is the term related to the initialization and is equal to 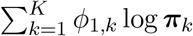.

#### M-step

The M-step of the Expectation-Maximization algorithm updates the GLM-HMM parameters by utilizing the posterior probabilities calculated during the E-step. The objective of the EM algorithm is to minimize the negative log-likelihood function, which is augmented with a prior on the GLM-HMM weights.

L-BFGS-B (Limited-memory Broyden–Fletcher–Goldfarb–Shanno with Bound constraints) is a popular optimization algorithm used to solve unconstrained and bound-constrained nonlinear optimization problems. It’s an extension of the BFGS algorithm, which is a quasi-Newton method for unconstrained optimization. L-BFGS-B is particularly useful when dealing with optimization problems where the variables have certain bounds or constraints. These constraints can be upper and lower bounds on the variables, and L-BFGS-B is designed to handle such constraints. So L-BFGS-B is a second-order optimization method that approximates the Hessian matrix, leading to faster convergence and accurate parameter estimates. Here, the EM algorithm employs the L-BFGS-B approach, which belongs to the class of Quasi-Newton optimization methods.

During the M-step, the EM algorithm maximizes the expected log-likelihood, computed through the forward-backward algorithm during the E-step. This optimization process aims to identify the best parameters for the GLM-HMM. Consequently, in each iteration of the EM algorithm, the initial state distribution ***π*** is updated to refine the model’s representation of the latent state dynamics.

By iteratively performing the E-step and M-step, the EM algorithm iteratively refines the parameter estimates until convergence is achieved, enabling the model to accurately capture the underlying dynamics and relationships in the data, leading to improved parameter estimation and enhanced model performance.

As mentioned, the EM algorithm calculates the expected log-likelihood, which is a concave function when considering the GLM weights, owing to the inclusion of a Bernoulli GLM in the set of functions that transform external inputs into probabilities of Hidden Markov Model emissions, as discussed in [25]. Since there is no closed-form solution for updating the GLM weights, the objective was to compute the global maximum of the expected log-likelihood. To achieve this, the scipy optimize function in Python [27] was used, which employs numerical optimization using the BFGS algorithm [28–31].

By applying the BFGS algorithm, the EM algorithm iteratively refines the GLM weights to maximize the expected log-likelihood, seeking the optimal parameters that best explain the observed data given the GLM-HMM. This numerical optimization approach enables a more computationally efficient determination of the parameter estimates, leading to improved model performance and accurate representation of the system’s dynamics.

In this study, the EM algorithm is employed with iterative steps until the difference between consecutive iterations falls below a specified tolerance, ensuring convergence to a stable solution. However, to further validate that the algorithm converges to the global optimum rather than a local optimum, a robust fitting procedure was adopted. Specifically, the EM algorithm’s weights were initialized multiple times (50 times), and for each initialization, the model was fitted. The consistency of the final results across these multiple runs demonstrated that the algorithm consistently converged to the accurate solution, representing the global optimum.

Moreover, to assess the uncertainty and estimate the posterior standard deviation of the model weights, the inverse Hessian of the optimized log-posterior was calculated. This computation was performed using Autograd, a powerful Python package and automatic differentiation library that simplifies and enhances gradient-based optimization.

#### Initialization

The initialization of the GLM-HMM weights followed a specific procedure. The observation weights for the GLM-HMM were initialized using a noisy version of a simple Bernoulli GLM. Initially, a 1-state GLM was fitted, and then the GLM-HMM was initialized with multiple states based on that basic GLM with added noise. This initialization approach provided a reasonable starting point for the GLM-HMM optimization process.

Regarding the transition weights, they were initialized with a vector of zeros, implying no initial knowledge about the transitions between states. Furthermore, to establish prior information for the initialization of the latent states, a uniform distribution prior was set. This choice of prior reflects a neutral assumption, where all states are considered equally probable at the outset.

Due to employing this initialization strategy and setting informative priors, we reached a systematic and principled starting point for the EM algorithm, facilitating more stable and reliable convergence to meaningful solutions during the subsequent parameter estimation and model fitting process.

#### 4.8.1 GLM-HMM fitting process

We present the results of applying the GLM-HMM independently to each mouse in Fig. 5. However, establishing a direct relationship between the obtained states across different animals poses challenges. To address this, we adopted a multi-step fitting technique. Initially, we combined the data from all mice into a unified dataset.

For the IBL dataset, we aggregated the data from all 37 mice. Employing Maximal Likelihood estimation and the EM algorithm, we analyzed this pooled data using a 1-state GLM-HMM, which represents a simpler GLM model. Subsequently, we used the obtained weights from this initial step as the initialization for the GLM weights during the fitting process of a *K*-state GLM-HMM. By doing so, we obtained a global fit by combining the dataset from all mice and fitting a *K*-state GLM-HMM. This approach allows us to assess the relationship between states across different animals and derive a unified representation of the latent states, providing valuable insights into the underlying dynamics observed across the entire dataset.

Following the global fit of the model, we proceeded with an individual fit, resulting in a distinct GLM-HMM fit for each animal. During this individual fit process, we initialized each animal’s model using the parameter values obtained from the global fit, which were derived from a model fitted to the pooled data from all mice. To identify the optimal initialization parameters, we conducted 50 initializations and compared the log-likelihood values for the training dataset, selecting the set of parameters that yielded the best fit. We then ran the EM algorithm until convergence for each animal’s model.

By employing this individual fitting approach with the best initialization parameters, there was no longer a need to permute the retrieved states of each animal to assign logically similar states to one another. Consequently, the recovered parameters are presented in Fig. 5, illustrating the distinct GLM-HMM fits for each mouse, facilitating an analysis of individual behavioral patterns and underlying dynamics.

In this analysis and fitting process, we incorporated Gaussian noise into the GLM weights enabling better discernment of the initialized states.

EM algorithms, like many optimization methods, can get stuck in local optima due to their reliance on initial parameter values. To overcome this issue, various strategies are employed, including multiple initializations, regularization, and exploration of the parameter space. These enhancements increase the chances of finding favorable solutions.

It’s crucial to recognize that achieving the global optimum of a likelihood function [32] is challenging, and different optimization techniques may be needed depending on the specific problem. Therefore, selecting the right optimization strategies tailored to the problem’s characteristics is essential for optimal parameter estimation and model fitting.

While EM algorithms can converge to local optima, they may not reach the global optimum. To address this, the model was fitted 50 times with 50 different initializations for each value of *K* to explore a broader range of solutions and identify the best fit.

As demonstrated here, the outlined initialization method exhibits sufficient stability and reliability, enabling the recovery of GLM-HMM parameters across a wide range of relevant parameter regimes in our analysis. This approach ensures robust estimation and fosters confidence in the model’s ability to accurately capture the underlying dynamics of the system even under diverse conditions.

#### 4.8.2 Fitting the psychometric curve

The psychometric function was derived by plotting the percentage of choices made by the animal against various stimulus values. In this paper, the psychometric curves illustrate the animal’s right-ward choice probability as a function of stimulus intensity. To fit the psychometric curve to the sigmoid function, maximum likelihood estimation was used, resulting in the following formulation:

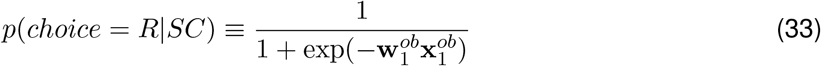

In which, the *SC* represents the stimulus contrast. To minimize the loss function, the study employed the Python package ‘optimize.minimize’ from the scipy library. This package provides numerical optimization tools that enable accurate minimization of the loss function, facilitating the estimation of the model parameters and the fitting of the psychometric curve.

### 4.9 K-fold Cross-validation for GLM-HMM

A cross-validation approach was employed to evaluate the model’s performance, involving random splitting of the data into 5 folds for training and testing. Approximately four-fifths of the data sessions were used for training the model, while the remaining randomly selected sessions were held out for testing the fitting performance. The sessions from all participating animals were evenly considered to ensure that mouse-to-mouse variability did not influence the cross-validation process.

This analysis revealed that a GLM-HMM with 3 to 5 latent states yields appropriate log-likelihood estimations. Following the fitting of the model with 1 to 5 states, we computed the log-likelihood of the test data. During the fitting procedure, the EM algorithm was executed on the training data sessions. Subsequently, in the testing stage, the likelihood of the remaining data was calculated based on the model parameters obtained from the training procedure. The forward-pass was performed only once during this stage, and the likelihood of the test data was obtained by summing over all states as 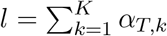 where *α*_*T,k*_ was computed solely on the held-out sessions. It is important to note that the GLM-HMM parameters were solely calculated during the training procedure and used for evaluating the model’s performance on unseen data during testing.

In our analysis, Fig. 2, we have calculated the log-likelihood using the described procedure which is the log of calculated likelihood and we call it *L* = log(*l*). In some results, we have used the unit term “bits per trial”. This *L* with this unit is acquired by calculating *L* of the test data on held out sessions as described and then subtracting log-likelihood of the identical dataset considering a Bernoulli model for observed data. This is a baseline model and its corresponding log-likelihood is shown by *L*_0_. Then we divided it by *T*_t_ log(2) in which *T*_t_ is test set size (the number of trials). The predictability of test sets can be greatly enhanced by even tiny amounts of log-likelihood, when expressed in bits per trial. To know more about how this can be explained, lets pay attention to a numerical example. A log-likelihood estimate of 0.02 bpt reveals that the test data is 984609.11 times more probable to have been made with the GLM-HMM than with a baseline model when the number of trials in the test data is 1000.”

In our analysis, we calculated the log-likelihood using the described procedure, denoted as *L* = log(*l*) and in some of the results, we used the unit term “bits per trial” to quantify the log-likelihood.

To obtain the log-likelihood in bits per trial (*L* bpt), we followed a specific process. Firstly, we computed *L* for the test data on held-out sessions, as described previously. Then, we determined the log-likelihood of the same dataset under the assumption of a Bernoulli model for observed data and multinomial GLM for transition data, which we refer to as the baseline model. We denoted the baseline model corresponding log-likelihood as *L*_0_. The difference between *L* and *L*_0_ represents the enhancement in log-likelihood due to the GLM-HMM performance. Finally, to express this enhancement per trial, we divided it by *T*_t_ log(2), where *T*_t_ is the size of the test set (the number of trials).

The resulting value in bits per trial provides a measure of predictability for the test set. Even minute increments in log-likelihood, when expressed in bits per trial, can lead to significant improvements in the predictability of the test data compared to the baseline model. To illustrate this, consider a numerical example where the log-likelihood difference estimate is 0.02 bpt. This implies that the test data is approximately 1048576 times more likely to have originated from the GLM-HMM compared to the baseline model when the test data comprises 1000 trials.

This representation in bits per trial, referred to as normalized log-likelihood, allows for a more intuitive understanding of the model’s performance and enables meaningful comparisons between different models and dataset. So we can write it as:

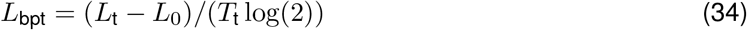

in which bpt is an abbreviation for bit per trial unit. Log-likelihood scores greater than 0 indicate no increase in estimation accuracy in comparison with the baseline model.

### 4.10 Dataset

In this framework, we analyzed data from 37 mice participating in the IBL decision-making task. each experimental trial encompasses the presentation of a sinusoidal grating, characterized by gradient values ranging from 0% to 100%. The data we used was during the stationary behavior phase after the training period. We considered both biased and unbiased blocks of the data. In the unbiased data, during the first 90 trials of each session, the probability of the stimulus appearing on the right or left side of the screen was equal. In contrast, for the biased data, the probability of the stimulus appearing on the left or right side was either 80% or 20% with random block lengths. The length of these biased blocks ranged from 20 to 100 trials. This dataset is publicly available on figshare at this link https://doi.org/10.6084/m9.figshare.11636748. We applied the GLM-HMM model to this data, and you can find more details in this paper [33].

## 5 Acknowledgement

We extend our sincere appreciation to Scott Linderman for his outstanding ‘Bayesian learning and state space modeling’ (ssm) package, from which our code in this paper is derived. We would like to acknowledge the feedback received from the International Brain Laboratory (IBL) and the Theory Working Group within IBL throughout the course of this research project. We are also grateful to Hannah Bayer and the IBL staff for their support. Special thanks go to our IBL board members, Liam Paninski, Alexander Pouget and Ilana B. Witten, for their constructive input on this manuscript. We express our gratitude to the members of the Pillow Lab especially Matthew Creamer.

## References

1. Ashwood, Z. C. et al. Mice alternate between discrete strategies during perceptual decision-making. Nature Neuroscience 25, 201–212 (2022).

2. Bolkan, S. S. et al. Opponent control of behavior by dorsomedial striatal pathways depends on task demands and internal state. Nature neuroscience 25, 345–357 (2022).

3. Ratcliff, R. & Rouder, J. N. Modeling response times for two-choice decisions. Psychological science 9, 347–356 (1998).

4. Ratcliff, R. & McKoon, G. The diffusion decision model: theory and data for two-choice decision tasks. Neural computation 20, 873–922 (2008).

5. Gold, J. I. & Shadlen, M. N. The neural basis of decision making. Annu. Rev. Neurosci. 30, 535–574 (2007).

6. Busse, L. et al. The Detection of Visual Contrast in the Behaving Mouse. en. Journal of Neuroscience 31, 11351–11361. ISSN: 0270-6474, 1529-2401. http://www.jneurosci.org/content/31/31/11351 (2019) x(Aug. 2011).

7. Brunton, B. W., Botvinick, M. M. & Brody, C. D. Rats and humans can optimally accumulate evidence for decision-making. Science 340, 95–98 (2013).

8. Pinto, L. et al. An Accumulation-of-Evidence Task Using Visual Pulses for Mice Navigating in Virtual Reality. English. Frontiers in Behavioral Neuroscience 12. Publisher: Frontiers. ISSN: 1662-5153. https://www.frontiersin.org/articles/10.3389/fnbeh.2018.00036/full?report=reader (2020) (2018).

9. Calhoun, A. J., Pillow, J. W. & Murthy, M. Unsupervised identification of the internal states that shape natural behavior. en. Nature Neuroscience 22. Number: 12 Publisher: Nature Publishing Group, 2040–2049. ISSN: 1546-1726. https://www.nature.com/articles/s41593-019-0533-x (2020) x(Dec. 2019).

10. Weilnhammer, V., Stuke, H., Eckert, A.-L., Standvoss, K. & Sterzer, P. Humans and mice fluctuate between external and internal modes of sensory processing. bioRxiv (2021).

11. Beron, C. C., Neufeld, S. Q., Linderman, S. W. & Sabatini, B. L. Mice exhibit stochastic and efficient action switching during probabilistic decision making. Proceedings of the National Academy of Sciences 119, e2113961119 (2022).

12. Le, N. M. et al. Mixture of Learning Strategies Underlies Rodent Behavior in Dynamic Foraging. bioRxiv, 2022–03 (2022).

13. Hulsey, D., Zumwalt, K., Mazzucato, L., McCormick, D. A. & Jaramillo, S. Decision-making dynamics are predicted by arousal and uninstructed movements. Cell Reports 43 (2024).

14. Baum, L. E., Petrie, T., Soules, G. & Weiss, N. A maximization technique occurring in the statistical analysis of probabilistic functions of Markov chains. The annals of mathematical statistics 41, 164–171 (1970).

15. Bengio, Y. & Frasconi, P. An input output HMM architecture. Advances in neural information processing systems 7 (1994).

16. Laboratory, T. I. B. et al. Standardized and reproducible measurement of decision-making in mice. Elife 10 (2021).

17. Burgess, C. P. et al. High-Yield Methods for Accurate Two-Alternative Visual Psychophysics in Head-Fixed Mice. eng. Cell Reports 20, 2513–2524. ISSN: 2211-1247 (Sept. 2017).

18. Berman, G. J., Choi, D. M., Bialek, W. & Shaevitz, J. W. Mapping the stereotyped behaviour of freely moving fruit flies. Journal of The Royal Society Interface 11, 20140672 (2014).

19. Wiltschko, A. B. et al. Mapping sub-second structure in mouse behavior. Neuron 88, 1121–1135 (2015).

20. Eyjolfsdottir, E., Branson, K., Yue, Y. & Perona, P. Learning recurrent representations for hierarchical behavior modeling. arXiv preprint arXiv:1611.00094 (2016).

21. Katsov, A. Y., Freifeld, L., Horowitz, M., Kuehn, S. & Clandinin, T. R. Dynamic structure of locomotor behavior in walking fruit flies. Elife 6, e26410 (2017).

22. Bishop, C. M. Pattern recognition and machine learning (springer, 2006).

23. Bolkan, S. S. et al. Opponent control of behavior by dorsomedial striatal pathways depends on task demands and internal state. Nature Neuroscience 25, 345–357. 10.1038/s41593-022-01021-9 (2022).

24. Dempster, A. P., Laird, N. M. & Rubin, D. B. Maximum likelihood from incomplete data via the EM algorithm. Journal of the royal statistical society: series B (methodological) 39, 1–22 (1977).

25. Escola, S., Fontanini, A., Katz, D. & Paninski, L. Hidden Markov models for the stimulus-response relationships of multistate neural systems. Neural computation 23, 1071–1132 (2011).

26. McLachlan, G. J. & Krishnan, T. The EM Algorithm and Extensions en. Google-Books-ID: NBawza-WoWa8C. ISBN: 978-0-470-19160-6 (John Wiley & Sons, Nov. 2007).

27. Virtanen, P. et al. SciPy 1.0: Fundamental Algorithms for Scientific Computing in Python. Nature Methods 17, 261–272 (2020).

28. Broyden, C. G. The convergence of a class of double-rank minimization algorithms 1. general considerations. IMA Journal of Applied Mathematics 6, 76–90 (1970).

29. Fletcher, R. A new approach to variable metric algorithms. The computer journal 13, 317–322 (1970).

30. Goldfarb, D. A family of variable-metric methods derived by variational means. en. Mathematics of Computation 24, 23–26. ISSN: 0025-5718, 1088-6842. https://www.ams.org/mcom/1970-24-109/S0025-5718-1970-0258249-6/ (2020) (1970).

31. Shanno, D. F. Conditioning of quasi-Newton methods for function minimization. Mathematics of computation 24, 647–656 (1970).

32. Salakhutdinov, R., Roweis, S. T. & Ghahramani, Z. Optimization with EM and expectation-conjugate-gradient in Proceedings of the 20th International Conference on Machine Learning (ICML-03) (2003), 672–679.

33. Aguillon, V. et al. A standardized and reproducible method to measure decision-making in mice. eLife (2020).

